# Identification of two bZIP transcription factors that regulate development of pavement and trichome cells in *Arabidopsis thaliana* by single-cell RNA-sequencing

**DOI:** 10.1101/2022.04.12.488054

**Authors:** Rui Wu, Zhixin Liu, Jiajing Wang, Weiqiang Li, Aizhi Qin, Xiaole Yu, Hao Liu, Chenxi Guo, Zihao Zhao, Yixin Zhang, Yaping Zhou, Susu Sun, Yumeng Liu, Mengke Hu, Jincheng Yang, Masood Jan, George Bawa, Jean-David Rochaix, Guoyong An, Luis Herrera-Estrella, Lam-Son Phan Tran, Xuwu Sun

**Affiliations:** State Key Laboratory of Crop Stress Adaptation and Improvement, State Key Laboratory of Cotton Biology, Key Laboratory of Plant Stress Biology, School of Life Sciences, Henan University, 85 Minglun Street, Kaifeng 475001, China; Departments of Molecular Biology and Plant Biology, University of Geneva, Geneva, 1211, Switzerland; Institute of Genomics for Crop Abiotic Stress Tolerance, Department of Plant and Soil Science, Texas Tech University, Lubbock, TX 79409, USA

**Author notes:** Correspondence: Contact: Dr. Xuwu Sun; Dr. Lam-Son Phan Tran; Dr. Luis Herrera-Estrella. These authors have contributed equally to this work (Rui Wu, Zhixin Liu, Jiajing Wang, Weiqiang Li, Aizhi Qin, Xiaole Yu). The author responsible for distribution of materials integral to the findings presented in this article in accordance with the policy described in the Instructions for Authors (https://academic.oup.com/plcell) is Xuwu Sun.

## Abstract

Epidermal cells are the main avenue for signal and material exchange between plants and the environment. Leaf epidermal cells primarily include pavement cells (PCs), guard cells, and trichomes cells (TCs), which differentiate from protodermal cells or meristemoids. The development and distribution of different epidermal cells are tightly regulated by a complex transcriptional regulatory network mediated by phytohormones, including jasmonic acid (JA), and transcription factors. Understanding how the fate of leaf epidermal cells is determined, however, is still largely unknown due to the diversity of cell types and the complexity of its regulation. Here, we characterized the transcriptional profiles of epidermal cells in 3-day-old true leaves of *Arabidopsis thaliana* using single-cell RNA-sequencing. We identified two genes encoding BASIC LEUCINE-ZIPPER (bZIP) transcription factors, namely the *bZIP25* and *bZIP53*, which are highly expressed in PCs and early-stage meristemoid cells. Densities of PCs and TCs were found to increase and decrease, respectively, in *bzip25* and *bzip53* mutants, compared with wild-type plants. This trend was more pronounced in the presence of JA, suggesting that these transcription factors regulate the development of TCs and PCs in response to JA.

**IN A NUTSHELL:** *Background:* Leaf epidermal cells, comprised of trichome cells (TCs), guard cells (GCs), and pavement cells (PCs), are responsible for exchanging materials and information between plants and the surrounding aerial environment. Many genes have been identified in *Arabidopsis thaliana* and confirmed to be involved in the initiation and differentiation of TCs and PCs. The fate determination of TCs and PCs is tightly regulated by positive and negative regulators at the cellular level. The precise underlying molecular mechanisms responsible for the fate determination of TCs and PCs, however, are still unclear at this time.

*Question:* What are the transcriptomic profiles of different leaf epidermal cell types? Can we dissect the genes that are specifically expressed in certain epidermal cell types? What kinds of transcription factors are involved in regulating the fate determination of TCs and PCs?

*Findings:* We performed single cell RNA-seq to investigate the transcriptomic profiles of different leaf epidermal cell types and identified differentially expressed genes in each cell type. We found that genes that are involved in jasmonic acid signaling are highly expressed in early-stage meristemoid (EM) cells which can act as the precursor of PCs and perhaps of TCs. To investigate the regulatory mechanisms underlying EM development, we identified the transcription factors (TFs) in EM cells and found that two bZIP TF genes, *bZIP25* and *bZIP53*, are highly expressed in EMs. Further analyses of these two genes using both loss-of-function and gain-of-function approaches indicated that bZIP25 and bZIP53 are functionally involved in promoting trichome formation but inhibit pavement cell development in response to jasmonic acid.

*Next steps:* Besides of bZIP25 and bZIP53, we also identified other key genes, for example *FES1B*, in leaf epidermal cells. Our next step will be to explore the regulation of other key genes involved in the fate determination of different cell types in leaf epidermis.

## Introduction

Epidermal cells are responsible for exchanging materials and information between the plants and the surrounding aerial environment (Pathuri et al., 2008). In leaves, epidermal cells can differentiate and produce trichomes, which are a specialized cell type that protect plants from adverse conditions including ultraviolet radiation and herbivore attack (Hauser, 2014). Thus, leaf epidermal cells are comprised of trichome cells (TCs), guard cells (GCs), and pavement cells (PCs) (Marks, 1997). Previous studies have systematically and comprehensively characterized the developmental dynamics of the transcriptomes of stomatal lineage cells (Liu et al., 2020). It is now important to examine the processes underlying the fates and development of PCs and TCs.

Growth and development of PCs in *Arabidopsis thaliana* mainly proceed through three stages. First, initial cells with different shapes begin to expand outward along the long axis of leaves to form outward elongated polygons. Then, the cells expand laterally along the edge of the adjacent cells, and subsequently extend irregularly to the side of the adjacent cells. Finally, the cells extend further outward, and the zigzagged protrusions are staggered with the narrow indentation of adjacent cells resulting in the formation of PC with different shapes (Fu et al., 2005). The irregular zigzagged protrusions of leaf epidermis are mainly regulated by the cytoskeleton (Xu et al., 2010). The dynamic arrangement of microtubules plays a role in the development of PCs (Eng et al., 2021). Microtubule-associated proteins KATANIN, IQ67 DOMAIN5 (IQD5), SPIRAL2 and CLASP are essential for morphogenesis of PCs (Ambrose et al., 2007; Lin et al., 2013; Wightman et al., 2013; Liang et al., 2018). Microfilaments mainly control the outward projection of the edge of epidermal cells (Armour et al., 2015). The Rho GTPase cascade signaling pathway is a foundation of the formation of PCs by activating microtubules and promoting their orderly arrangement which consequently leads to morphological changes of leaf epidermal cells (Lin et al., 2013).

TCs are cells that originate on the epidermis of aerial organs and serve as an excellent model for the study of differentiation in plants at the cellular level (Marks, 1997). TCs are regularly spaced and rarely appear adjacent to each other, suggesting that TC spacing is a tightly regulated process (Lloyd et al., 1994; Larkin et al., 1996; Schnittger et al., 1999; Esch et al., 2004; Zhao et al., 2008; Hilscher et al., 2009; Balkunde et al., 2010; Pesch and Hulskamp, 2011; Grebe, 2012; Yanagisawa et al., 2015). More than 40 genes involved in the initiation and differentiation of TCs have been identified in Arabidopsis (*Arabidopsis thaliana*) (Hulskamp et al., 1994; Marks et al., 2009). For example, mutations in several transcription factor (TF)-encoding genes like *GLABROUS1* (*GL1*), *GLABRA2* (*GL2*), *TRANSPARENT TESTA GLABRA1* (*TTG1*), or both *GLABRA3* (*GL3*) and *ENHANCER OF GLABRA3* (*EGL3*) result in the loss of TCs (Koornneef, 1981; Herman and Marks, 1989; Marks and Feldmann, 1989; Oppenheimer et al., 1991; Hulskamp et al., 1994; Larkin et al., 1994; Rerie et al., 1994; Szymanski et al., 1998; Payne et al., 1999; Walker et al., 1999; Payne et al., 2000; Esch et al., 2003; Zhang et al., 2003; Kirik et al., 2005). Notably, both positive and negative activity of regulatory TFs are required for the development of TCs (Ishida et al., 2008; Pesch and Hulskamp, 2009). Positive regulators in Arabidopsis include members of several TF families, such as MYB, basic helix–loop–helix (bHLH), WDR, and C2H2 zinc finger families. R2R3-MYB TFs include GL1 and its paralog MYB23 (Herman and Marks, 1989; Marks and Feldmann, 1989; Kirik et al., 2001; Kirik et al., 2005; Ishida et al., 2008; Marks et al., 2009). GL1 and MYB23 have been reported to be functionally equivalent during TC initiation but not during the TC branching process (Kirik et al., 2005). The bHLH family members include GL3 and its homolog EGL3 that are involved in TC development in a partially redundant manner (Payne et al., 2000; Morohashi et al., 2007; Hao et al., 2019). TC development is activated by TTG1, a protein containing a WD40 repeat, a highly conserved motif consisting of approximately 40–43 amino acids, often ending with Trp-Asp (W-D) residues (Walker et al., 1999; Zhang et al., 2003). Both GL1 and TTG1 control the same process in TC development (Herman and Marks, 1989; Marks and Feldmann, 1989; Oppenheimer et al., 1991; Larkin et al., 1994; Walker et al., 1999; Payne et al., 2000; Kirik et al., 2005). Negative regulators of TC development include at least seven MYB proteins: CAPRICE (CPC), TRIPTYCHON (TRY), ENHANCER OF TRY AND CPC1 (ETC1), ETC2, ETC3, TRICHOMELESS1 (TCL1), and TCL2 (Wada et al., 1997; Kirik et al., 2004; Kirik et al., 2004; Zhu et al., 2009; Gan et al., 2011; Tian et al., 2017). These negative regulators exhibit partially redundant roles in the initiation and differentiation of TCs. This fact is evidenced, for example, by functional studies of three genes: *CPC, ETC2*, and *ETC3*, in which the *cpc etc2 etc3* triple mutant exhibited an increased density of TCs compared with the single mutants (Wada et al., 1997; Kirik et al., 2004; Wang et al., 2008; Hilscher et al., 2009; Wester et al., 2009; Zhu et al., 2009). Interestingly, one study found that *ETC3* is highly expressed in young stomatal cells, and that its expression is under the control of SPEECHLESS (SPCH) which is highly expressed in early-stage meristemoid (EM) cells (Adrian et al., 2015). These results suggest that SPCH may activate genes that promote trichome differentiation, and that EM cells may act as the precursor cells for TC production.

Several other TFs, phytohormones, and cell developmental factors can also affect TC development (Walker et al., 2000; Breuer et al., 2009; Yoshida et al., 2009; Wen et al., 2018; Vadde et al., 2019). The TF AtMYC1 was identified as a direct target of both GL1 and GL3 in Arabidopsis (Pesch et al., 2013). Furthermore, a TEOSINTE BRANCHED1, CYCLOIDIA, and PROLIFERATING CELL NUCLEAR ANTIGEN FACTOR1/2 (TCP) TF family member, namely TCP4, was reported to affect TC fate by directly binding to the promoters of *TCL1* and *TCL2*, which in turn repress *GL2* (Efroni et al., 2008). The positive regulators of TC initiation GL1 and GL2 are significantly up-regulated in *TCP4* loss-of-function mutants and down-regulated in gain-of-function mutants (Vadde et al., 2019). Previous studies have also indicated that wounding and jasmonate (JA) significantly promote TC initiation (Traw and Bergelson, 2003; Li et al., 2004; Boughton et al., 2005; Qi et al., 2011; Tian et al., 2016; Yan et al., 2017). Qi et al., (2011) found that JA ZIM-domain (JAZ) repressor proteins interact with bHLH (Transparent Testa8, GL3, and EGL3) and R2R3-MYB TFs (e.g., MYB75 and GLABRA1), which are the essential components of WD-repeat/bHLH/MYB transcriptional complexes (Qi et al., 2011). JAZ proteins are substrates of the CORONATINE INSENSITIVE1 (COI1)–based SCF^COI1^ E3 ligase complex (Gupta et al., 2021). Upon JA binding with COI1, COI1 recruits JAZ proteins to the SCF^COI1^ E3 complex for ubiquitination and degradation through the 26S proteasome (Chini et al., 2007). Subsequently, the bHLH and MYB components of WD-repeat/bHLH/MYB complexes are released and the development of TCs is activated (Qi et al., 2011). Regarding the TC development, two proteins, SIAMESE (SIM) and STICHEL (STI), play a fundamental role in the regulation of the endoreduplication of nuclear DNA (Walker et al., 2000; Ilgenfritz et al., 2003; Churchman et al., 2006).

Several transcriptomic studies of TC development have been conducted to explore the regulatory processes responsible for the development of TCs and to identify new regulatory factors associated with TC development (Marks et al., 2008; Marks et al., 2009; Wang et al., 2009; Chen et al., 2014; Yang et al., 2015; Akhtar et al., 2017). These initial studies have characterized the transcriptome of TCs and identified several marker genes associated with the regulation of TC development and function. Transcriptome analysis of TCs alone, however, cannot dissect the regulation of the fate determination of TCs at the gene expression level. This is because the fate determination of TCs is also affected by fate determination factors and the developmental status of PCs that are adjacent to TCs, which can potentially differentiate into TCs (Grebe, 2012). The fate determination of TCs and PCs is tightly regulated by positive and negative regulators at the cellular level. The precise underlying molecular mechanisms responsible for the fate determination of TCs and PCs, however, are still unclear. Single-cell RNA-sequencing (scRNA-seq) technology allows the analysis of transcriptional profiles of different types of cells and to identify genes that are specifically expressed at different developmental and morphogenetic stages (Zhang et al., 2019; Liu et al., 2020; Wendrich et al., 2020; Kim et al., 2021; Liu et al., 2021; Serrano-Ron et al., 2021; Liu et al., 2022). Therefore, we conducted a scRNA-seq analysis of 3-day-old true leaves of Arabidopsis wild-type (WT) to elucidate the mechanisms that regulate the fate and development of TCs and PCs. Our study identified a group of novel marker genes for PCs and TCs, and discovered the new roles of two BASIC LEUCINE-ZIPPER (bZIP) TFs in the regulation of fate determination and differentiation of PCs and TCs through the comparative analysis of WT and single and double mutants of *bZIP25* and *bZIP53* genes.

## Results

### Single-cell transcriptional profiles of leaf epidermal cells unravels different cell types and gene expression signatures

We subjected protoplasts of 3-day-old true leaves of Arabidopsis to scRNA-seq analysis to identify cell type-specific changes in gene expression that occur during epidermal cell fate determination at a single-cell resolution (Figure 1A-D). Protoplasts were filtered through a 40 μm cell strainer and cell viability was assessed by phenol blue staining. A total of 18,000 cells were subsequently used to generate the libraries that were sequenced (Figure 1B and C). After stringent cell filtration, high-quality transcriptomes of 15,773 individual cells were retained for subsequent analyses (Figure 1D). A total of 512,130,798 reads were obtained after processing the sequencing data, with an average of 32,468 reads and 2,118 genes identified per cell. The percentage of reads mapped to the genome was 93%. We then performed t-distributed stochastic neighbor embedding (tSNE) dimensionality analysis of the scRNA-seq data. Supplemental Figure S1A and B illustrate the tSNE projection plots of cells colored by unique molecular identifier (UMI) counts and automated clustering, respectively.

**Figure 1.**
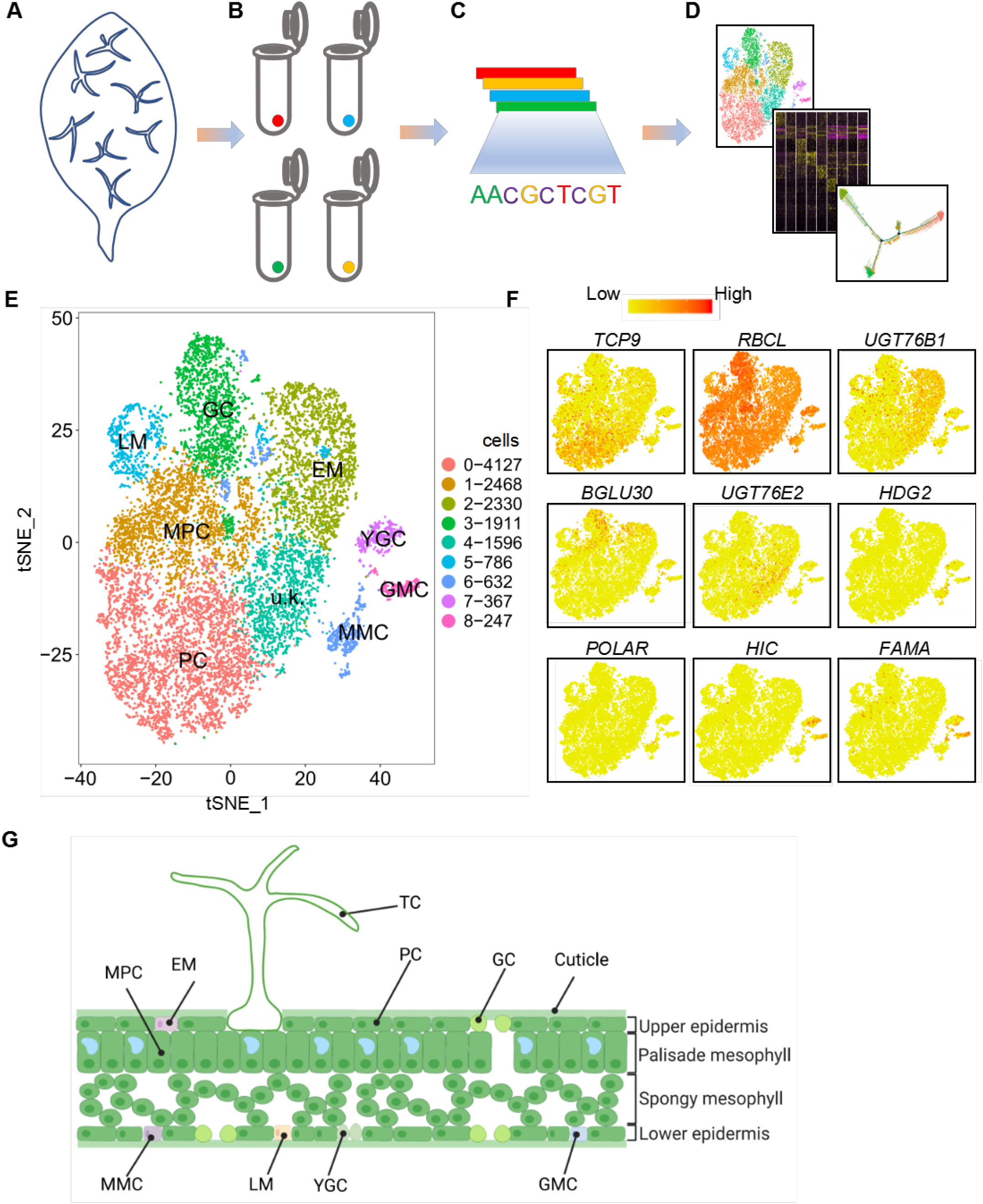
Distinct cell subpopulations with transcriptional signatures determined by single-cell RNA-sequencing analysis of epidermal cells of true leaves. (A-D) Illustration of the scheme used for young leaves (A), cell isolation (B), sequencing (C), and downstream analysis (D). E, t-distributed stochastic neighbor embedding (tSNE) plot reveals cellular heterogeneity with 9 distinct clusters of cells identified and color-coded. General identity of each cell cluster is defined in the corresponding cluster. F, Feature plots of expression distribution for selected marker genes. Expression levels for each cell are color-coded and overlaid onto the tSNE plot. G, Illustration of a leaf section with the different cell types. TC, trichome cell; EM, early-stage meristemoid; GC, guard cell; PC, pavement cell; LM, late-stage meristemoid; YCG, young guard cell; MPC, mesophyll cell; GMC, guard mother cell; MMC, meristemoid mother cell; u.k., unknown.

The sequencing saturation satisfied the requirement of 10×genomics (Supplemental Figure S1C). The median number of genes per cell (using TAIR10 as the reference genome) also met the requirement for data analysis (Supplemental Figure S1D). We then analyzed the scRNA-seq data by principal component analysis (PCA). Supplemental Figure S1E shows the distribution of the percent of mitochondrial gene sequences (percent. mito) on a PCA plot. Supplemental Figure S1F and G display the UMI distribution (nUMI) and number of nuclear-encoded genes (nGene) on the PCA plot. After removing mitochondrial and chloroplast transcripts, a total of 14,464 cells were used for the subsequent analysis (Supplemental Figure S1H). Subsequently, tSNE analysis was carried out on the selected cells. As shown in Figure 1E, 9 cell clusters were identified as being independently distributed on the tSNE plot. We also visualized cell clusters using the uniform manifold approximation and projection (UMAP) algorithm on our scRNA-seq data (Supplemental Figure S2). The UMAP analysis produced similar cell clusters as those in the tSNE analysis (Supplemental Figure S2A and S2B). We identified differentially expressed genes (DEGs) in the different cell types (Supplemental Table S1 and Table S2). The expression patterns of the top 10 marker genes in each cell type are shown in a heatmap plot (Figure 2A and Supplemental Figure S2C). The violin plots and feature plots of representative marker genes in each cell type are shown in Figure 2B and C and Supplemental Figure S2D.

**Figure 2.**
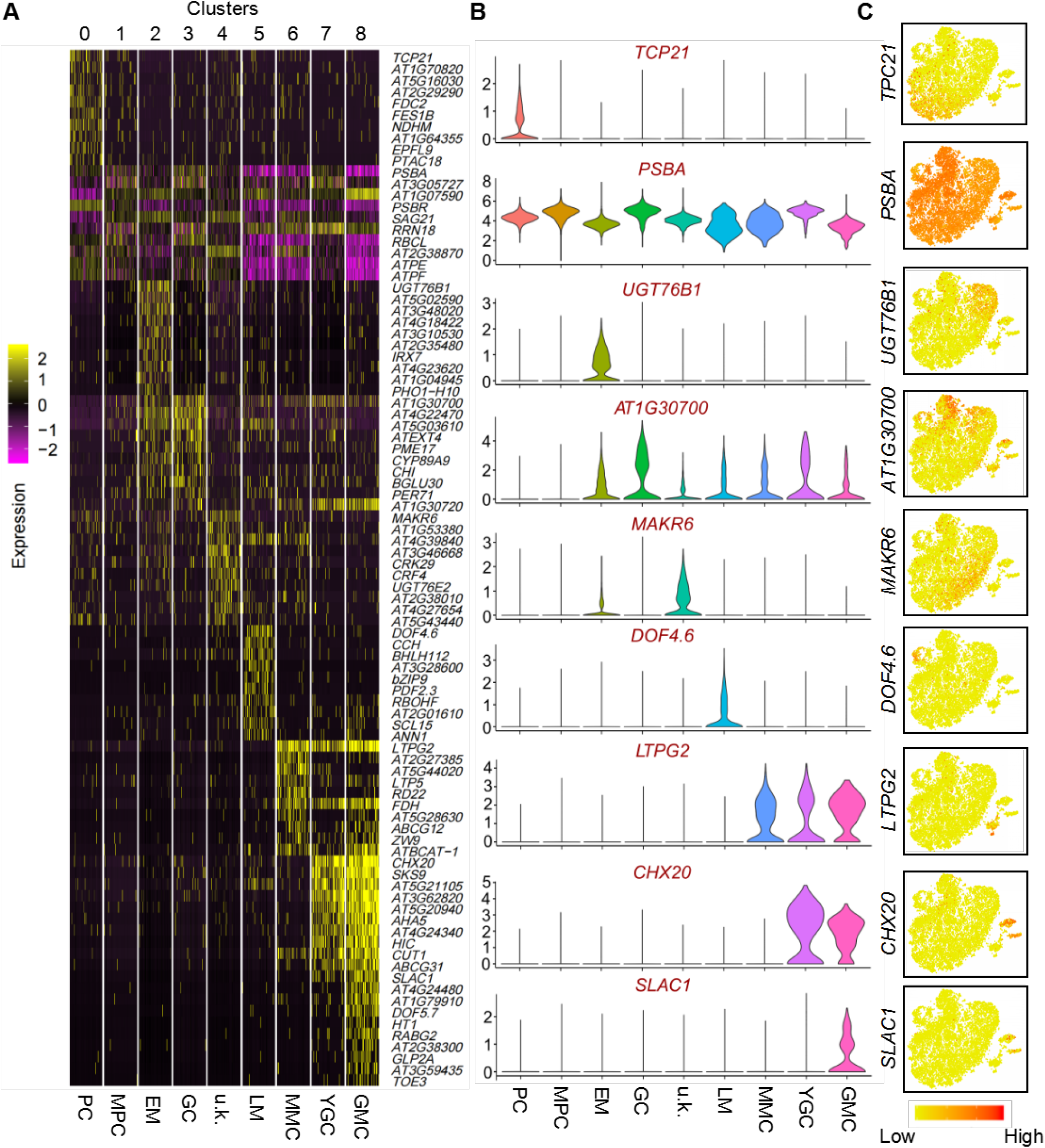
Identification of novel marker genes for each cluster. A, Heatmap of differentially expressed genes (DEGs). The top 5 genes and their relative expression levels in all sequenced cells are shown for each cluster. The color ranges from purple to yellow and represents the expression value of the marker genes from low to high. B, Violin plots of selected novel marker genes for each cluster. C, Feature plots of the expression distribution of selected novel marker genes. Expression levels for each cell are color-coded and superimposed on the tSNE plot. EM, early-stage meristemoid; GC, guard cell; PC, pavement cell; LM, late-stage meristemoid; YCG, young guard cell; MPC, mesophyll cell; GMC, guard mother cell; MMC, meristemoid mother cell; u.k., unknown.

We then determined the cell type of the identified cell clusters using well-defined cell type-specific marker genes for epidermal cells. As shown in Figures 1F and 2A-C, the epidermal marker gene for PCs, *TCP21*, was chiefly expressed in cluster 0; the marker gene for mesophyll cells (MPCs), *RIBULOSE BISPHOSPHATE CARBOXYLASE LARGE CHAIN* (*RBCL*), was primarily distributed in cluster 1; the marker gene for EMs, *UDP-DEPENDENT GLYCOSYLTRANSFERASE 76B1* (*UGT76B1*), was mainly expressed in cluster 2; the marker gene for GCs, *BETA-GLUCOSIDASE* (*BGLU30*), was predominantly distributed in cluster 3; the late-stage meristemoid cells (LMs) marker genes, *DNA BINDING WITH ONE FINGER 4*.*6* (*DOF4*.*6*) and *bZIP9*, were mainly enriched in cluster 5; the meristemoid mother cell (MMC) marker gene, *HOMEODOMAIN GLABROUS 2* (*HDG2*), was chiefly expressed in cluster 6; the young guard cells (YGCs) marker gene, *HIGH CARBON DIOXIDE* (*HIC*), was mainly expressed in cluster 7; the guard mother cell (GMC) marker genes, *FAMA* and *DOF5*.*7*, were mostly expressed in cluster 8. However, the marker gene for TCs, *GL2* (Szymanski et al., 1998), was unfortunately not detected in our scRNA-seq data, perhaps because the size of TCs was too large to pass through the cell strainer. Thus, no known marker genes were identified to be expressed in cluster 4. Collectively, our results indicate that cluster 0 belongs to PCs; cluster 1 belongs to MPCs; cluster 2 belongs to EMs; cluster 3 belongs to GCs; cluster 4 belongs to unknown (u.k.) cells; cluster 5 belongs to LMs; cluster 6 belongs to MMCs; cluster 7 belongs to YGCs; cluster 8 belongs to GMCs. Notably, the expression of some marker genes of the JA signal transduction pathway, such as *ACYL-COA OXIDASE 1* (*ACX1*) (Peng et al., 2019), *ABNORMAL INFLORESCENCE MERISTEM* (*AIM1*) (Delker et al., 2007), *BLADE ON PETIOLE1* (*BOP1*) (Canet et al., 2012), *CORONATINE INSENSITIVE 1* (*COI1*) (Xie et al., 1998; Thines et al., 2007), *CONSTITUTIVE EXPRESSION OF PR GENES 5* (*CPR5*) (Clarke et al., 2001), *CULLIN 1* (*CUL1*) (Quint et al., 2005); *JASMONATE-ZIM-DOMAIN PROTEIN 10* (*JAZ10*) (Chung and Howe, 2009), *JASMONATE-INDUCED OXYGENASE2* (*JAO2*), *JASMONATE-INDUCED OXYGENASE3* (*JAO3*) (Caarls et al., 2017), *PRODUCTION OF ANTHOCYANIN PIGMENT 1* (*PAP1*) (Bali et al., 2019), *RADICAL-INDUCED CELL DEATH1* (*RCD1*) (Overmyer et al., 2000), and *RIBONUCLEASE 1* (*RNS1*) (LeBrasseur et al., 2002) was also significant in cluster 2, suggesting that they may function in EMs (Supplemental Figure S3).

### Selection and characterization of newly identified marker genes in PCs and Ems

Gene ontology (GO) analysis was then performed to identify the potential biological function of DEGs in each cell cluster (Supplemental Table S3). As shown in Supplemental Figure S4, GO terms in MPCs and PCs were generally very similar and different from the other cell types. GO terms in the u.k., LM, GMC, and EM clusters were comparable, suggesting that these genes are involved in similar biological processes in these different cell types. GO terms for MPCs were predominantly related to photosynthesis, consistent with the functions of MPCs (Supplemental Figure S4). Considering the high similarity in GO terms in clusters 3 and 7, we propose that cluster 3 belongs to GCs (Supplemental Figure S4). We could not identify TCs by the expression of the TC marker gene *GL2* because the size of TCs was too large to pass through the cell strainer. In our previous study, we found that some marker genes were detected in several cell types but at different levels of expression (Liu et al., 2020). The top 10 marker genes for each of the studied cell types other than TCs were specifically expressed in the corresponding cell types, except for the markers of MPCs, GMCs, and GCs. Some marker genes of PCs, such as *FERREDOXIN C 2* (*FDC2*), *FES1B, AT2G29290*, and *EPIDERMAL PATTERNING FACTOR LIKE-9* (*EPFL9*), were also enriched in MPCs and GCs (Figure 2A).

Transgenic plants expressing yellow fluorescent protein (YFP) fusion proteins of some of the representative genes were generated to determine the cellular localization of the proteins encoded by the selected marker genes. The expression of YFP was detected in PCs for TCP21, FDC2, NADH DEHYDROGENASE-LIKE COMPLEX M (NDHM), AT1G70820, AT2G29290, and AT5G02590 (Supplemental Figure S5). YFP signals for FDC2 and NDHM were also detected in GCs. The localization of AT5G02590 and PECTIN METHYLESTERASE 17 (PME17) in PCs and EMs could not be distinguished from each other. YFP signals for bZIP9 were detected in GCs, although its expression was mostly in LMs (Figure 2A and Supplemental Figure S5). In general, the expression patterns of the selected genes were consistent with the results presented in Figure 2A. We also constructed promoter-driven *GUS* reporter gene vectors for some of the representative genes to analyze the tissue specificity of gene expression. *AT2G29290, FES1B, TCP21, PLASTID TRANSCRIPTIONALLY ACTIVE 18* (*PTAC18*), *FDC2*, and *AT1G64355* were selected as marker genes for PCs, while *AT1G04950* and *EUKARYOTIC RELEASE FACTOR 1-2* (*ERF1-2*) were selected as marker genes for EMs, and the corresponding transgenic plants were produced. GUS staining analysis revealed that the selected genes are expressed in the leaves of seedlings (Figure 3). Some genes, such as *TCP21, FDC2, ERF1-2*, and *AT1G04950*, can also expressed in roots (Figure 3), suggesting these genes may also play an important function in root epidermal cells.

**Figure 3.**
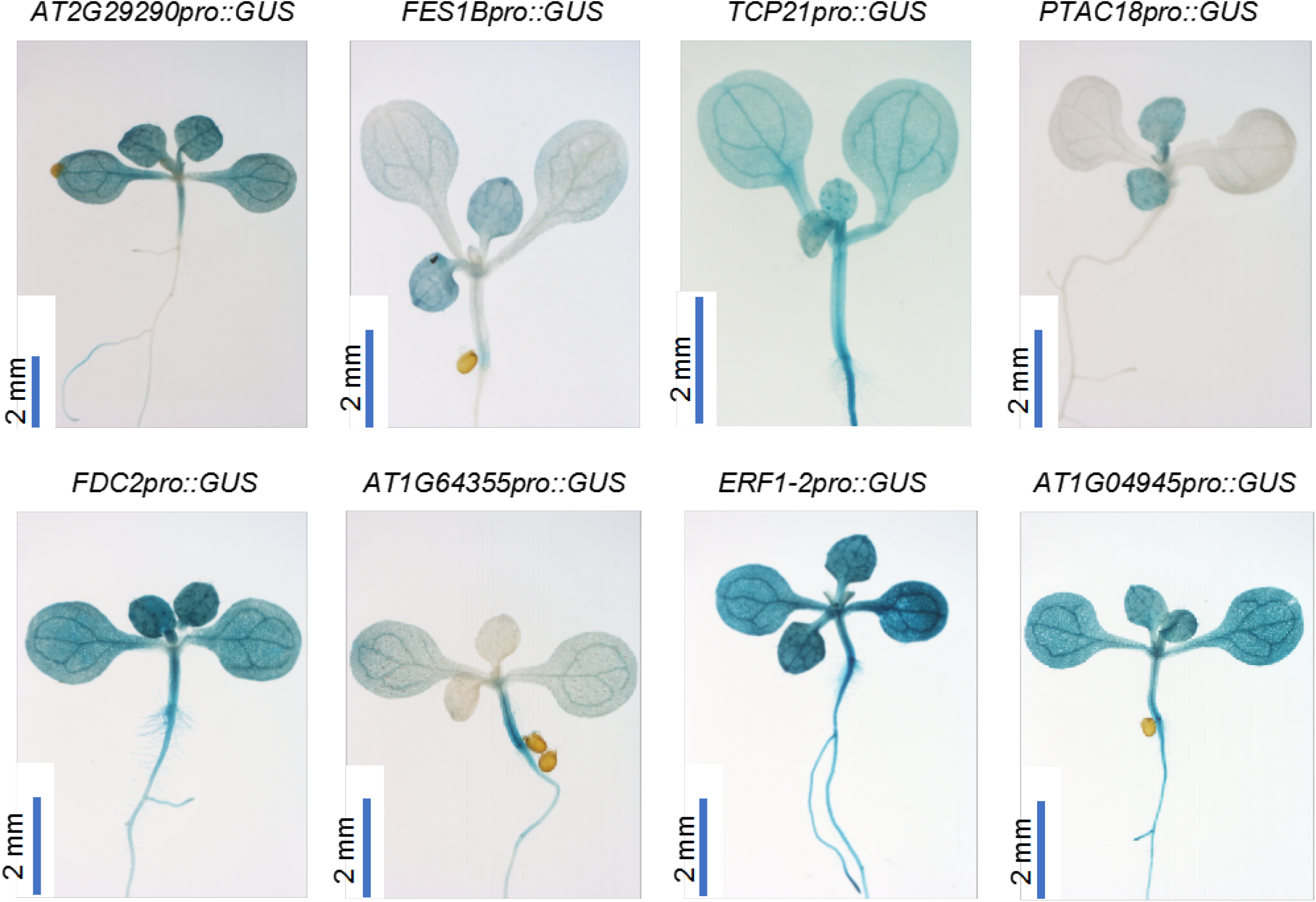
Expression of selected marker genes in different tissues. Transgenic plants expressing the *GUS* reporter gene driven by the promoters of the selected marker genes were generated to analyze their expression patterns. GUS signals were detected by staining. Scale bar, 2 mm is shown as a blue line.

Transgenic plants were also successfully produced that overexpressed several selected newly identified PC (e.g., *TCP21, AT1G70820, AT2G29290, FDC2, FES1B, NDHM, EPFL9*, and *PTAC18*) and EM (e.g., *AT5G02590, AT3G48020, AT4G18422, AT3G10530*, and *AT4G23620*) marker genes to determine their potential roles in the regulation of PC and TC development. Results indicated that, compared with WT plants, overexpression of *TCP21, FDC2, AT4G18422*, and *AT4G23620* resulted in a significant decrease in the density of TCs, while overexpression of *FES1B, NDHM*, and *AT3G10530* enhanced the density of TCs (Figure 4A and C). PC density was significantly lower, relative to WT plants, in seedlings of *35S::FES1B, 35S::PTAC18, 35S::FDC2*, and *35S::AT3G48020* transgenic plants, but significantly higher in *35S::EPFL9* and *35S::AT4G23620* transgenic seedlings (Figure 4B and D). Leaf area in *35S::TCP21* plants was also significantly smaller than in WT plants (Figure 4E). These results suggest that the selected PC and EM marker genes may be involved in the development of PCs and TCs.

**Figure 4.**
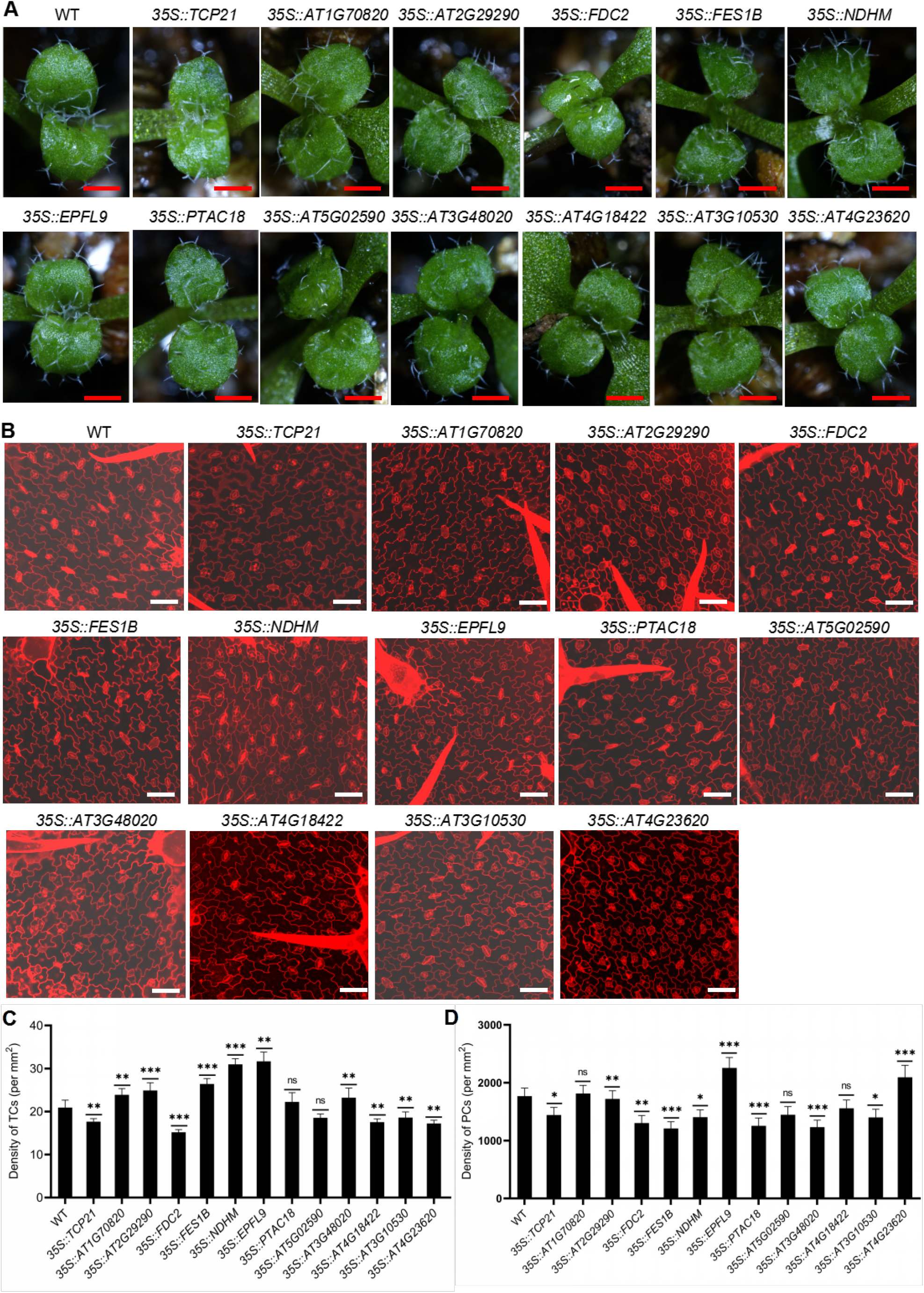
Characterization of the potential roles of selected marker genes for pavement cells (PCs) and trichome cells (TCs). A, Detection of the developmental status of trichomes in transgenic lines and wild-type (WT) seedlings. Scale bar (0.5 mm) is shown as a red line with a white background. B, Analysis of the developmental status of PCs in WT and transgenic lines. Samples were stained with propidium iodide, after which PCs were detected using a laser confocal microscope. Scale bar (50 μm) is shown as a white line. C, TC density in the upper epidermis of two 3-day-old true leaves of WT and transgenic seedlings. D, PC density in the upper epidermis of 3-day-old true leaves of WT and transgenic seedlings. Data represent a mean ± SD (*n* = 3). Asterisks indicate a significant difference between transgenic and WT plants as determined using a Student’s *t*-test. \**P* < 0.05, \*\**p* < 0.01, and \*\*\**p* < 0.001. ns, non-significant.

### Pseudo-time trajectory analysis of the spatiotemporal dynamics of epidermal cell differentiation

Arabidopsis leaf development is a strictly regulated process that ensures that almost all leaves have similar spatial morphological characteristics at the same developmental stage (Byrne et al., 2001; Fleming, 2005; Bar and Ori, 2014). The spatiotemporal regulation of leaf development is closely related to that of cell development (Kalve et al., 2014; Lu et al., 2014). Therefore, understanding the spatiotemporal regulation pattern of cell development is important for understanding leaf development. Taking this into consideration, we performed a pseudo-temporal ordering of cells (pseudo-time) on the scRNA-seq data using Monocle 2 (Trapnell et al., 2014) to reconstruct the developmental trajectory during differentiation. The resulting pseudo-time path has two nodes and three branches (Figure 5A), and different cell clusters are arranged relatively clearly at different branch sites of the pseudo-time path (Figure 5B). A heatmap analysis based on pseudo-time results was then constructed to characterize the spatiotemporal dynamic patterns of the top 10 genes of each cluster. As shown in Figure 5C, the heatmap of several representative genes from each cluster indicated a positive correlation between their expression dynamics and their cell distribution on the developmental trajectory. For example, *UGT76B1* and *PEROXIDASE 71* (*PER71*) are maximally expressed in the pre-branch of the pseudo-time trajectory, while *TPC21* and *EPFL9* are mostly expressed in the late stage of cell fate 1 (Figure 5C). Marker genes of stomatal lineage cells, such as *HIC* and *DOF5*.*7*, have their highest expression levels in the early stage of cell development, while these genes are down-regulated following the developmental direction of cell fate 1 and cell fate 2 (Figure 5C). These results indicate that genes expressed in different cell types have a specific spatiotemporal pattern on the pseudo-time trajectory.

**Figure 5.**
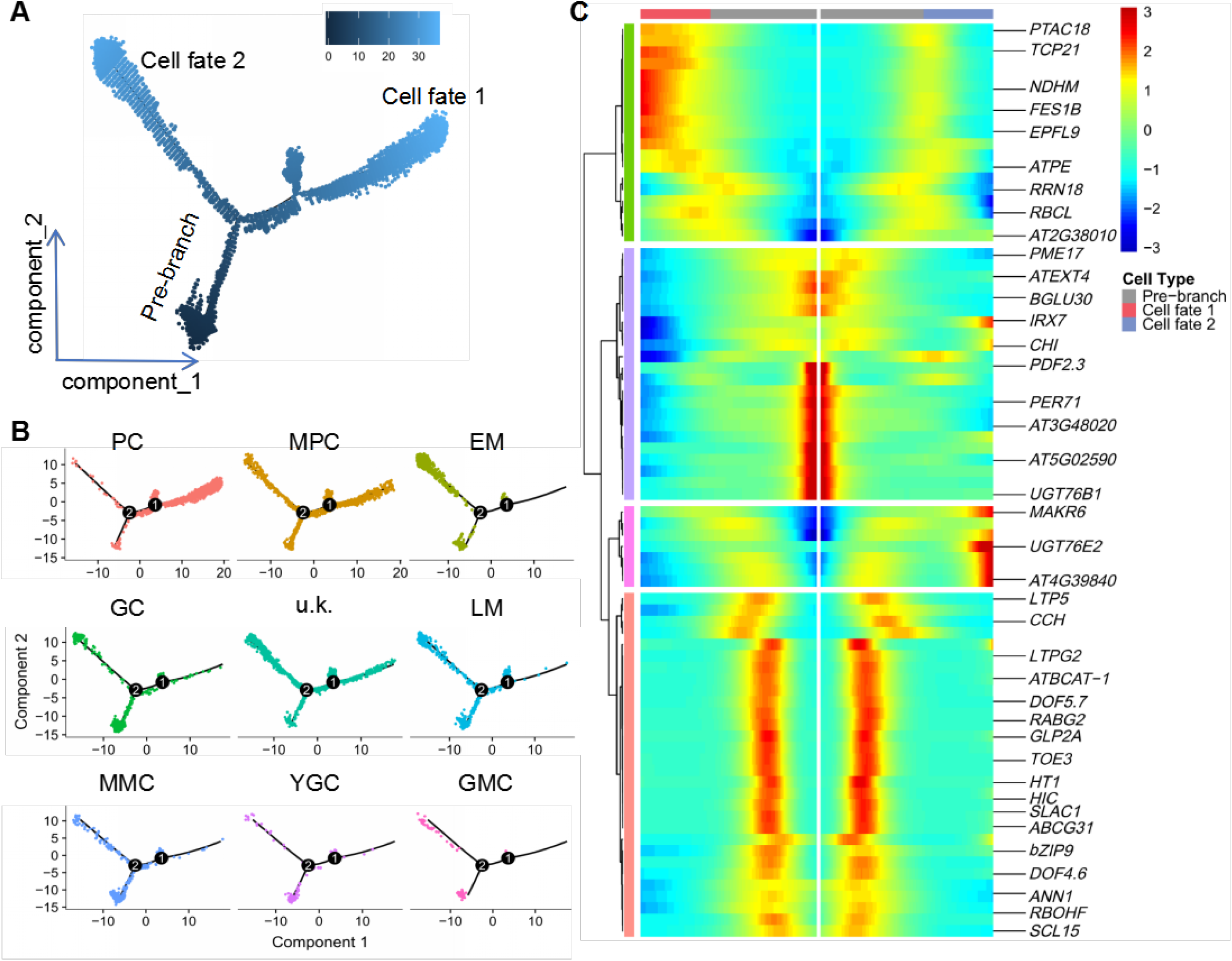
Pseudo-time analysis reveals putative differentiation trajectories of different cell types. A, Distribution of cells of each cluster on the pseudo-time trajectory. B, Distribution of cells of each cell type on the pseudo-time trajectory. C, Clustering and expression kinetics of the top 10 genes in all clusters along with a pseudo-time progression. EM, early-stage meristemoid; GC, guard cell; PC, pavement cell; LM, late-stage meristemoid; YCG, young guard cell; MPC, mesophyll cell; GMC, guard mother cell; MMC, meristemoid mother cell; u.k., unknown.

### Analysis of the effects of JA on the development of TCs and PCs

Since JA signaling marker genes are expressed in EMs (Supplemental Figure S3), it is possible that JA could be involved in the regulation of EM differentiation. It has been proposed that EMs give rise to both PCs and TCs (Adrian et al., 2015), and JA plays important roles in regulating the development of TCs (Yan et al., 2017). Therefore, to explore this possibility, we first analyzed the process of TC differentiation in WT seedlings in the presence of JA. Results indicated that the number of TCs significantly increased in the presence of 20 μM JA (Supplemental Figure S6). Higher concentrations of JA (> 40 μM) inhibit leaf growth, although the density of TCs gradually increases with the increasing JA dose (Supplemental Figure S6B and C). We then analyzed the effects of JA on the development of PCs, and found that the density of PCs decreased along with the increasing JA concentrations (0 to 40 μM) (Supplemental Figure S7).

### bZIP TFs are involved in regulating the fate of PCs and TCs

TFs play important roles in regulating the development of all kinds of cells. For example, SPCH, FAMA, MUTE, and BASIC PENTACYSTEINE 6 (BPC6) TFs are essential for the development of GCs (Liu et al., 2020). In our search for potential regulators of PCs and TCs, we identified two TF-encoding genes, *bZIP25* and *bZIP53*, that were predominantly expressed in EMs and PCs (Figure 6A and B), raising the possibility that they may be involved in regulating the fate and differentiation of these cells. GUS expression can be detected in the true leaves of *bZIP25pro::GUS* transgenic plants (Supplemental Figure S8A). Notably, YFP signals in bZIP25-YFP plants were highly detected in the nuclei of PCs (Supplemental Figure S8B). We then examined the corresponding T-DNA insertion mutants *bzip25* and *bzip53* obtained from the Arabidopsis Biological Resource Center (ABRC) to investigate the potential roles of bZIP25 and bZIP53 in the regulation of PC and TC development. The developmental states of TCs in leaves of the single mutant *bzip25* and *bzip53* seedlings are shown in Figure 6C. Results indicated that TC densities in *bzip25* and *bzip53* seedlings were lower than in WT plants with and without JA treatment, but the response to JA was decreased in the mutants as compared with the WT (Figure 6E). In contrast, the analysis of PCs showed that PC density in *bzip25* and *bzip53* mutant seedlings was higher than in the WT with and without JA-treatment (Figure 6F). Consistently, a greater number of TCs was observed in *35S::bZIP25* and *35S::bZIP53* plants, while the number of PCs in *35S::bZIP25* and *35S::bZIP53* plants was lower than the WT (Figure 6C-F). These results indicate that bZIP25 and bZIP53 play a positive role in determining the density of TCs and a negative role in the density of PCs

**Figure 6.**
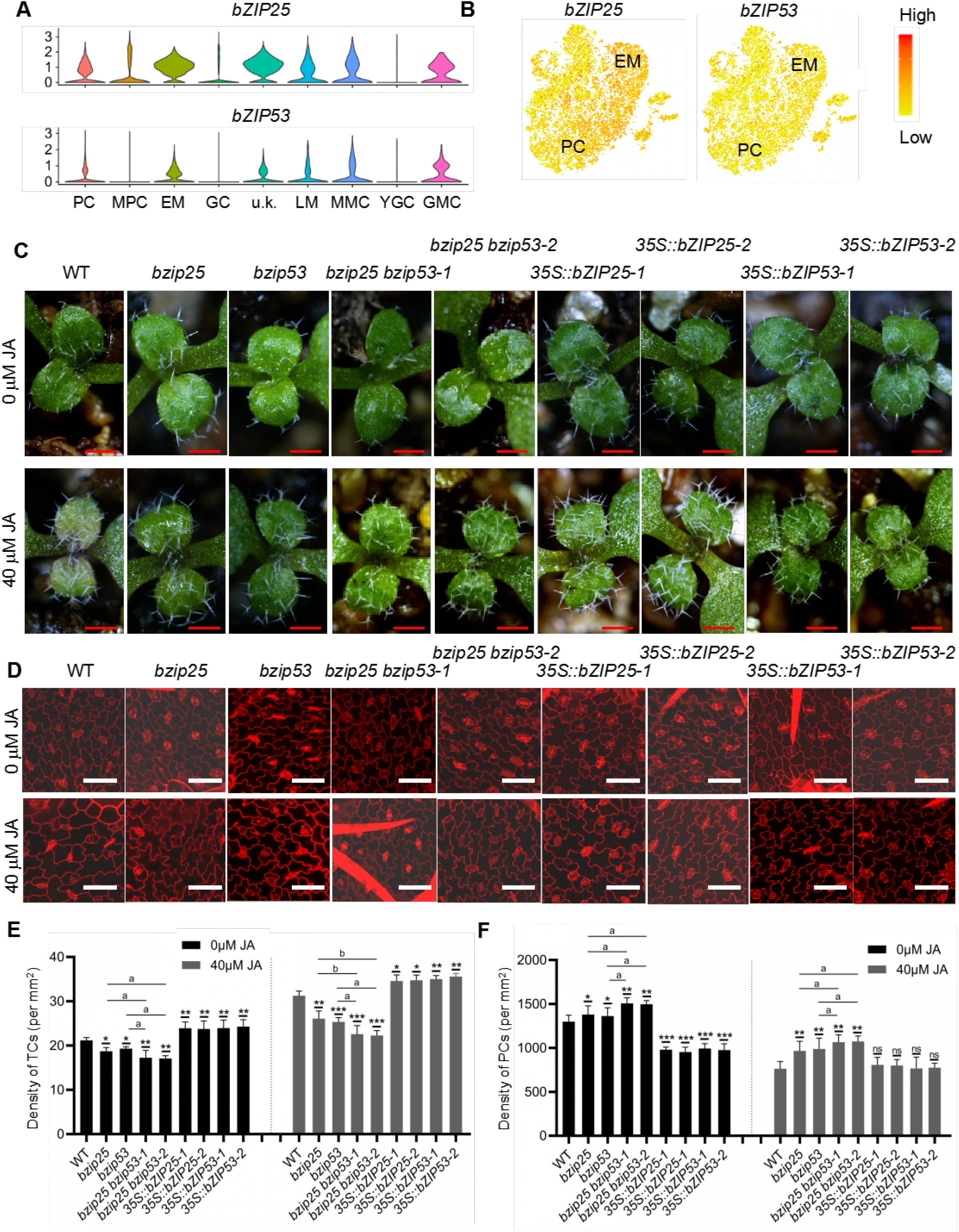
bZIP25 and bZIP53 positively regulate trichome cell (TC) development and negatively regulate pavement cell (PC) development. A, Violin plots showing the expression of *bZIP25* and *bZIP53* in EMs and PCs. B, Feature plots showing the expression of *bZIP25* and *bZIP53* in EMs and PCs. C, Representative photographs of the upper epidermis of 3-day-old true leaves of WT, *bzip25, bzip53, bzip25 bzip53, 35S::bZIP25* and *35S::bZIP53* plants subjected to 0 (control) and 40 μM jasmonic acid (JA) treatments. Bar, 500 μm. D, Representative photographs of PCs in the upper epidermis of 3-day-old true leaves of WT, *bzip25, bzip53, bzip25 bzip53, 35S::bZIP25* and *35S::bZIP53* plants subjected to 0 (control) and 40 μM jasmonic acid (JA) treatments. The samples were treated by propidium iodide (PI) staining to show the cell wall. Bar, 50 μm. (E-F) Density of TCs (E) and density of PCs (F) in the upper epidermis of two 3-day-old true leaves of WT, *bzip25, bzip53, bzip25 bzip53, 35S::bZIP25* and *35S::bZIP53* plants subjected to 0 and 40 μM JA treatments. Data represent a mean ± SD (*n* = 3). Asterisks indicate a significant difference between mutant and WT, and between overexpression lines and WT as determined using a Student’s *t*-test. \**P* < 0.05, \*\**p* < 0.01, and \*\*\**p* < 0.001. ns, non-significant. Letters indicate a significant difference between single mutant and double mutant as determined using a Student’s *t*-test. ^a^ *P* < 0.05, ^b^ *P* < 0.01.

Next, to test whether *bZIP25* and *bZIP53* function in the same regulatory pathway of epidermal cell development, we generated the double mutants *bzip25 bzip53-1* and *bzip25 bzip53-2* using CRISPR/Cas9 technology (Figure 6C and Supplemental Figure S9). Under control conditions (no JA), the number of TCs was lower in the double mutants, relative to the single mutants and WT plants, while the number of PCs was greater (Figure 6D and F). The effects of JA on TC and PC development were weak in the double mutants, relative to the single mutants and WT plants (Figure 6A-D). These results collectively suggest that *bZIP25* and *bZIP53* might play additive or partially redundant roles in regulating the fate and differentiation of PCs and TCs (Figure 7).

**Figure 7.**
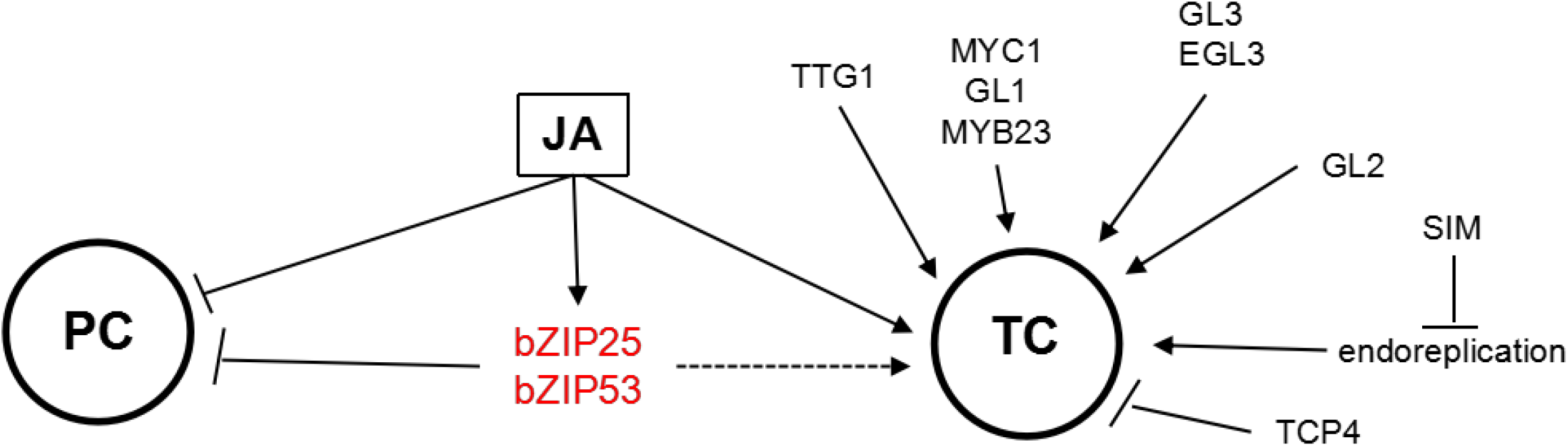
Model of the transcription factor network, including bZIP25 and bZIP53, in regulating TC and PC development. The fate of a TC is regulated by a series of critical transcription factors, including TTG1, GL2, MYC1, GL3, SIM, and TCPs, as well as others. JA positively promotes the development of TC but inhibits the differentiation of PC. bZIP25 and bZIP53, in response to JA, negatively regulate the fate of a PC and may positively promote the differentiation of a TC.

## Discussion

### Identification of marker genes in cell clusters obtained by scRNA-seq of young leaves

Utilizing scRNA-seq technology, we constructed the global landscape of the transcriptomes of young epidermal cell types in leaves. Unlike cotyledons, leaf epidermal cell types are more complex with the most striking feature being the development of TCs. The fate determination and differentiation of TCs are tightly regulated by both internal factors, such as hormones, and external cues, such as invading pests and pathogens (Ishida et al., 2008). The transcriptome of TCs has been extensively characterized but not at the single-cell level (Marks et al., 2008; Marks et al., 2009). Also, for true leaves, no reported studies on epidermal cells at single-cell resolution are available. TCs are differentiated from protodermal cells or EMs (Adrian et al., 2015). Therefore, a comprehensive study of the transcriptomes of different epidermal cell types in true leaves will enable us to identify the potential key regulators of the differentiation and development of different cell types in the leaf epidermis. Because EMs, PCs and TCs of true leaves have certain interaction in differentiation and development (Adrian et al., 2015), we can identify the regulatory factors regulating TCs by analyzing the key regulatory factors in EMs and PCs. In the scRNA-seq data obtained in this study, we did not identify the cell type in which the well-known TC marker gene *GL2* is specifically expressed (Figure 1). One possible explanation is that the size of TCs is too large and filtered out during the process of cell filtration used to prepare protoplast for scRNA-seq. Therefore, in this work we mainly focused on the characterization of the transcriptomes of EMs and PCs. We identified several genes that specifically expressed in EMs and PCs (Figure 2 and Supplemental Figure S2). To verify the cell types identified in the present study, we generated *GUS* reporters constructs for representative marker genes in PCs and EMs (Figure 3). Analysis of the expression pattern of the *GUS* reporter constructs indicated that the marker genes for PCs (including *AT2G29290, FES1B, TCP21, PTAC18*, and *FDC2*) and EMs (including *AT1G04950, AT4G18422, bZIP25*, and *ERF1-2*) are highly expressed in true leaves (Figure 3). Since EM cells and PCs are distributed among the entire upper epidermal layer of true leaves, *GUS* expression, which was controlled by the promoter of the marker genes expressed in EMs and PCs, appears to occur in all epidermal cells (Figure 3). We also generated transgenic plants expressing YFP fusion proteins of some of the representative marker genes to detect their cell expression pattern. As predicted, the expression of PC marker genes, such as *AT2G29290, FDC2, NDHM, AT1G70820*, and *TCP21*, were detected in PCs (Supplemental Figure S5). The expression of the EM marker genes, *AT5G02590* and *PME17*, however, was also observed in PCs (Supplemental Figure S5), which is consistent with the fact that PCs develop from EM cells (Adrian et al., 2015) (Figure 1G). In our study, *bZIP9* expression was detected in GCs (Supplemental Figure S5), however, a recent study has reported that *bZIP9* is strongly expressed in phloem parenchyma cells (Kim et al., 2021). *NDHM* and *FDC2* were also found to express in GCs (Supplemental Figure S5). The specific expression of the examined marker genes in PCs and EM cells suggests that these genes may be involved in mediating the development of these two cell types (Figure 3). Analysis of the developmental status of TCs and PCs in seedlings of transgenic plants overexpressing selected newly identified marker genes revealed that *TCP21, FDC2, AT5G02590, AT4G18422*, and *AT4G23620* negatively affect TC development, while *FES1B, NDHM*, and *EPFL9* positively affect TC development (Figure 4). Our results also demonstrated that *FES1B* negatively regulates the development of PCs, while *EPFL9* and *AT4G23620* regulate PC development in a positive manner (Figure 4). At present, the distinction between PCs and EM cells is difficult due to the inability to define specific marker genes. Our results provide important data that can be used for identifying PCs and EM cells in future scRNA-seq studies of epidermal cell development.

### Dissection of the spatiotemporal patterns of the transcriptomes of epidermal cells in true leaves

GCs, PCs, EM cells, and TCs are the main cell types present in the upper epidermis of leaves of *A. thaliana*. PCs, EM cells, and GCs differentiate from MMCs. According to the distribution of cells in the constructed pseudo-time trajectories, MMCs mainly appear at the initial stage, while EM cells and GCs are distributed over the later stages of pseudo-time trajectories (Figure 5B). This is consistent with the viewpoint that GCs and EMs differentiate from MMCs (Liu et al., 2020). Formation of TCs was highly similar to that of EMs in regard to developmental regulation (Adrian et al., 2015). Therefore, the results of the pseudo-time trajectory of EMs also support the evidence indicating that EMs or TCs are differentiated from MMCs. Pseudo-time heatmap analysis of the top 10 genes further confirmed this premise (Figure 5C). Our results indicate that the analysis of the spatiotemporal patterns of gene expression in specific types of cells significantly contributes to the understanding of their development.

### bZIP TFs are involved in regulating the fate and development of EMs and TCs in response to JA signaling

Identification of key TFs in specific cell types can assist in the identification of important regulatory factors involved in the fate determination and development of specific cell types. *bZIP25* and *bZIP53* were identified in our analysis of TF-encoding genes with increased expression in PCs and EMs (Figure 6A and B). Previous studies have shown that JA promotes the development of TCs (Yan et al., 2017), but inhibits the development of leaves (Noir et al., 2013). Our results demonstrated that high concentrations of JA inhibited leaf growth, but that TCs density gradually increased as the applied dose of JA increased (Supplemental Figure S6B and C). Further analysis of the effects of JA on the developmental status of both PCs and TCs in seedlings of the *bzip25* and *bzip53* single and double mutants revealed that *bZIP25* and *bZIP53* may have additive or partially redundant functions in the regulation of development of PCs and TCs (Figure 7). In summary, our results provide new insights into the mechanisms underlying the highly complex yet orderly orchestrated process of epidermal cell development. These findings provide a basis for the further study of novel regulators of specific cell types in the epidermis of leaves.

## Materials and methods

### Screening and verification of mutants

Wild-type (WT) and *A. thaliana* (Col-0 ecotype) were used in the scRNA-seq experiments. Seeds were sterilized in 5% sodium hypochlorite and germinated on vertical, half-strength Murashige and Skoog (1/2 MS) plates. T-DNA insertion mutants were obtained from the Arabidopsis Biological Resource Center (ABRC) (Supplemental Table 4). Mutant lines homozygous for the T-DNA insertion were identified by PCR analysis using gene-specific and T-DNA-specific primers (Supplemental Table 5 and Supplemental Figure S10). All mutants and WT plants were grown in a climate chamber at 22°C and 100 µmol photons m^−2^ s^−1^ under a 14-h light/10-h dark regime. In the experiments designed to examine the effect of JA on the TC development, 3-day-old seedlings were treated by spraying methyl jasmonate (392707, Millipore Sigma, St Louis, MO, USA). The seedlings were then placed in a sealed transparent plastic container that allowed them to continue to grow for a defined period of time. The developmental status of TCs was photographically documented.

### Constructs for plant transformation

*YFP-fusion expression constructs* - full-length cDNA fragments of marker genes were PCR-amplified using the primer pairs described in Supplemental Table 5. The resulting PCR products were purified and cloned into pDNOR201 by BP Clonase reactions (GATEWAY Cloning; Invitrogen, Waltham, MA, USA) according to the manufacturer’s instructions to generate the pDONR-cDNA vectors. The resulting plasmids were then recombined into pB7YWG2.0 using LR Clonase reactions to generate the final constructs.

*GUS reporter constructs* - the upstream 2,000-bp fragments of marker genes were PCR-amplified using the primer pairs described in Supplemental Table 5. The resulting PCR products were purified and cloned into pDNOR201 by BP Clonase reactions according to the manufacturer’s instructions to generate the pDONR-cDNA vectors. The resulting plasmids were recombined into pBGWFS7 using LR Clonase reactions to generate the final constructs. The resulting reporter constructs were then used to detect the expression of GUS under the control of the promoters of the different marker genes.

### Plant transformation

YFP-fusion expression constructs and reporter constructs were transformed into *Agrobacterium tumefaciens* strain GV3101 via electroporation. *A. tumefaciens* containing the different constructs were introduced into WT plants. The resulting T1 transgenic plants containing YFP-fusion expression constructs and reporter constructs were selected using BASTA as described previously (Sun et al., 2016). Homozygous transgenic plants were used in all experiments.

### Sample collection and protoplast preparation

Three-day-old true leaves were harvested and used to isolate protoplasts as previously described with slight modifications to adjust for the use of young leaf tissues (Yoo et al., 2007; Liu et al., 2020).

### ScRNA-seq library preparation

ScRNA-seq libraries were prepared using a Chromium Single Cell 3’ Gel Beads-in-emulsion (GEM) Library & Gel Bead Kit v3 according to the manufacturer’s instructions (10× Genomics, California, USA).

### ScRNA-seq data preprocessing

The rew data were processed as previously described (Liu et al., 2020). After the critical filtering process, 14,464 out of 15,773 cells were retained for downstream analysis. The median value of the mapping rate was 66.8%, and the median number of genes detected in each cell was 2,118. Library size normalization was performed in Seurat on the filtered matrix to obtain normalized counts.

### Clustering analysis of scRNA-seq data

Genes with the greatest variable expression amongst single cells were identified using the method described by (Macosko et al., 2015). The tSNE analysis, UMAP analysis, and DEG identification were performed as previously described (Liu et al., 2020; Liu et al., 2022).

### Pseudo-time and trajectory analysis

Pseudo-time trajectory analysis of single-cell transcriptomes was conducted using Monocle 2 (Trapnell et al., 2014) as previously described (Liu et al., 2020).

### Bulk and scRNA-seq correlation analysis

Bulk and scRNA-seq correlation analysis was performed as described (Rheaume et al., 2018). Differential expression was analyzed with a *t*-test. The *t*-test function was used to test the gene expression value in scRNA-seq and bulk, and a significant *P*-value was obtained. The difference multiple of log_2_ (FC) was calculated as follows: log_2_ ([mean gene expression value in scRNA-seq] + 0.001) / ((mean gene expression value in bulk) + 0.001). Finally, genes with a significant difference in expression were identified based on a *p*-value < 0.05 and | log_2_ (FC) | > 1.

### RNA-seq analysis

Three-day-old true leaves were harvested for extraction of total RNA using a mirVana miRNA Isolation Kit (Ambion, Waltham, MA, USA) following the manufacturer’s protocol. Samples with an RNA Integrity Number (RIN) ≥ 7 were subjected to subsequent RNA-seq analysis. Libraries were constructed using a TruSeq Stranded mRNA LT Sample Prep Kit (Illumina, San Diego, CA, USA) according to the manufacturer’s instructions. Libraries were sequenced on an Illumina sequencing platform (HiSeqTM 2500 or Illumina HiSeq X Ten), and 125-bp/150-bp paired-end reads were generated.

### GUS staining and histological analysis

Histochemical GUS staining was performed with a G3061 GUS staining Kit (Solarbio Co., Beijing, China) according to the manufacturer’s instructions as previously described (Liu et al., 2022).

### Microscopy

Seedlings were stained with 10 g mL^−1^ propidium iodide (PI) (P4170, Sigma, St Louis, MO, USA) for 1 min prior to imaging. PI staining was used to stain the cell wall of epidermal cells. Fluorescence in roots was detected using a Zeiss LSM980 confocal laser scanning microscope (Zeiss, Oberkochen, Germany). The PI signal was visualized at 610 to 630 nm wavelengths. YFP was observed at 510 to 530 nm wavelengths. Images and GFP intensities were processed using Zeiss Confocal Software.

### GO enrichment analysis

GO enrichment pathway analyses for the DEGs were conducted in Metascape (http://metascape.org/) (Zhou et al., 2019).

### Accession numbers

The accession numbers for some of the selected genes are as follows: *AT5G08330* (*TCP21*), *AT3G53800* (*FES1B*), *AT3G54620* (*bZIP25*), *AT3G58750* (*CSY2*), *AT3G62420* (*bZIP53*), *AT1G32550* (*FDC2*), *AT3G11340* (*UGT76B1*), *AT2G28110* (*FRA8*/*IRX7*), *AT4G12970* (*STOMAGEN*/*EPFL9*), *AT1G12920* (*ERF1-2*), *AT2G42790* (*CSY3*), *AT4G37925* (*NDHM*), *AT2G32180* (*PTAC18*), *AT1G69480* (*PHO1-H10*), *AT1G70820, AT5G16030, AT2G29290, AT1G64355, AT5G02590, AT3G48020, AT4G18422, AT3G10530, AT2G35480, AT4G23620* and *AT1G04945*. ScRNA-seq data are available at the following web addresses: (https://dataview.ncbi.nlm.nih.gov/?search=SUB6947465; https://www.ncbi.nlm.nih.gov).

## Supplemental data

The following materials are available in the online version of this article.

**Supplemental Figure S1**. Analysis of the single-cell RNA-sequencing (scRNA-seq) raw data.

**Supplemental Figure S2**. Comparative analysis of the cell clusters identified with different methods of dimensionality reduction.

**Supplemental Figure S3**. The expression pattern of jasmonic acid (JA) marker genes.

**Supplemental Figure S4**. GO and KEGG analysis of the differentially expressed genes (DEGs) in each of the cell clusters.

**Supplemental Figure S5**. Analysis of the expression of marker genes.

**Supplemental Figure S6**. Jasmonic acid (JA) promotes the development of TCs.

**Supplemental Figure S7**. Jasmonic acid (JA) inhibits the development of PCs.

**Supplemental Figure S8**. Analysis of the expression of bZIP25.

**Supplemental Figure S9**. Analysis of the DNA sequence in *bzip25* and *bzip53* seedlings after DNA editing.

**Supplemental Figure S10**. Verification of the T-DNA insertion in mutants.

**Supplemental Table 1**. All_DEGs_for_all_clsuters_tSNE.

**Supplemental Table 2**. All_DEGs_for_all_clusters_UMAP.

**Supplemental Table 3**. GO analysis of differentially expressed genes (DEGs).

**Supplemental Table 4**. List of mutant lines used in the study.

**Supplemental Table 5**. List of oligonucleotide primer pairs used in the study.

## Acknowledgments

We are grateful to ABRC for the *Arabidopsis* seeds. This research was supported by the National Natural Science Foundation of China (31670233).

## Author contributions

Conceptualization of the project:X.S. Experimental design: X.S and Z.L. Performance of some specific experiments: R.W., Z.L. J.W., A.Q., X.Y., H.L., Z.Z., Yixin Zhang, C.G., Yaping Zhou, M.J., G.B., S.S., Y.L., M.H., and J.Y. Data analysis: W.L., J.W., Y.Z., G.A. and J.R. Manuscript drafting: S.X. Contribution to the editing and proofreading of the manuscript draft: J.R., L.E., and L.T. All authors have read and approved the final manuscript.

## Data Availability Statement

All data supporting the findings of this study are available within the paper and the supplementary data published online.

